# The FAIR-Device - a non-lethal and generalist semi-automatic Malaise trap for insect biodiversity monitoring: Proof of concept

**DOI:** 10.1101/2024.03.22.586299

**Authors:** Juan A. Chiavassa, Martin Kraft, Patrick Noack, Simon Walther, Ameli Kirse, Christoph Scherber

**Author notes:** /.

## Abstract

Field monitoring plays a crucial role in understanding insect dynamics within ecosystems. It facilitates pest distribution assessment, control measure evaluation, and prediction of pest outbreaks. Additionally, it provides important information on bioindicators with which the state of biodiversity and ecological integrity in specific habitats and ecosystems can be accurately assessed. However, traditional monitoring systems can present various difficulties, leading to a limited temporal and spatial resolution of the obtained information. Despite recent advancements in automatic insect monitoring traps, also called e-traps, most of these systems focus exclusively on studying agricultural pests, rendering them unsuitable for monitoring diverse insect populations. To address this issue, we introduce the Field Automatic Insect Recognition (FAIR)-Device, a novel non-lethal field tool that relies on semi-automatic image capture and species identification using artificial intelligence via the iNaturalist platform. Our objective was to develop an automatic, cost-effective, and non-specific monitoring solution capable of providing high-resolution data for assessing insect diversity. During a 26-day proof-of-concept evaluation, the FAIR-Device recorded 24.8 GB of video, identifying 431 individuals from 9 orders, 50 families, and 69 genera. While improvements are possible, our device demonstrated potential as a cost-effective, non-lethal tool for monitoring insect biodiversity. Looking ahead, we envision new monitoring systems such as e-traps as valuable tools for real-time insect monitoring, offering unprecedented insights for ecological research and agricultural practices.

## Introduction

A large number of recent studies have documented or at least suggested potentially severe declines in biodiversity at various spatiotemporal scales, with many studies suggesting declines in insect taxa such as flying insects in general, wild bee pollinators, or butterflies [1, 2]. At the same time, we lack consistent and efficient approaches to non-invasively monitor insect populations [3]. Many current monitoring exercises deploy Malaise traps (e.g. the European Malaise trap program) that catch hundreds or even thousands of insect species per week, which can be problematic, especially in protected areas. It is practically impossible to manually sort the masses of insects collected by such traps, and even metabarcoding approaches take weeks or months for data to emerge after lab processing. Developing real-time monitoring systems and non-lethal traps is thus a priority for many biodiversity monitoring endeavors worldwide.

In recent years, advancements in sensors, processors, network infrastructure, and information management systems-such as artificial intelligence (AI) and big data-have paved the way for a multitude of new monitoring devices that provide unprecedented access to environmental information [4–7]. Among these innovations, electronic traps (e-traps) have emerged as a stand-out solution. Designed to replace traditional insect monitoring methods, these traps employ various technologies to gather data on insect populations, behavior, and diversity. These traps provide real-time, wirelessly transmitted data that undergo continuous analysis. Leveraging advanced sensor technologies, e-traps are at the forefront of monitoring solutions, particularly those that include image acquisition and processing [3, 4, 8, 9].

AI technologies play a crucial role in this context, with recent breakthroughs in deep learning, computer vision, and image processing achieving remarkable accuracy. As a result, manual in-situ observations and routine laboratory sample processing can now be surpassed [3, 8]. However, most e-traps developed thus far have primarily focused on studying agricultural pests, tailoring their trap and bait designs to specific species or a limited range of insects [10]. This specialization renders them less suitable for monitoring highly heterogeneous insect communities and poses challenges for conservation science due to their often lethal nature. In summary, while technological progress has revolutionized insect monitoring, there remains a critical need for non-lethal, non-specific solutions capable of comprehensively assessing insect diversity across ecosystems.

Recently developed non-invasive systems that utilize cameras and AI algorithms for insect detection and identification provide an exception to lethal automated monitoring devices. For instance, the Insect Detect DIY camera trap is designed to video-monitor a colored surface through tracking and recognition algorithms [11, 12]. This surface is optimized to attract mainly hoverflies (*Syrphidae*) and *Hymenoptera*. The trap is based on cost-effective off-the-shelf hardware components combined with open-source software. Light traps that use computer vision have also been presented [13, 14]. The Automated Moth Trap (AMT), for example, uses a combination of light and a camera to attract and detect nocturnal moths. The video-captured moths are then identified, tracked, counted, and classified using image processing algorithms. These non-lethal insect monitoring devices address ethical concerns while providing reliable and accurate data. However, they are designed to detect specific types of insects and therefore cannot be considered non-specific traps.

The use of smart mobile devices as biodiversity monitoring tools has also gained popularity in recent times, due to the development of smartphone apps for taxonomic identification. These apps often offer AI image identification functionalities for the taxonomic classification of species, and many of them are constantly improving as algorithms and computing power advance [15]. Two of the most popular taxonomic identification applications are iNaturalist (https://www.inaturalist.org) and ObsIdentify (https://observation.org), with millions of downloads on Google Play [16, 17] and in Applés App Store [18, 19]. Both apps allow users to record, identify, and share observations of wild organisms with other users of the platform community. The community is actively involved in curating the observations by adding identifications to observations made by others, which helps to improve the accuracy of the data. The platforms also provide a way for users to connect with others, build their knowledge, and contribute to science.

Biodiversity assessments using mobile apps are usually considered non-invasive, greatly expanding the possibilities of conservation monitoring in near-real time (especially compared to lethal insect traps). However, the main purpose of such portals is the opportunistic recording of species occurrence and not structured ecological monitoring [3]. Additionally, this type of active monitoring necessarily requires the presence of the observer in situ, making it a labor-intensive task.

Here, we introduce the Field Automatic Insect Recognizer-Device (FAIR-Device), a generalist and non-lethal field stationary device based on semi-automatic image capture and species identification by artificial vision through the iNaturalist platform. Our objective was to develop a cost-effective monitoring system that allows harmlessly and with high temporal resolution to identify and count insects on the field. Although many e-traps have been presented in recent years to automatically count and/or identify insects, we here aim at a truly non-specific monitoring device that directly builds upon the traditional Malaise trapping systems.

We hypothesize that

i. a system such as the FAIR-Device will allow non-lethal monitoring of a wide range of taxa of different species, and
ii. the processing of the acquired images with the iNaturalist AI system will greatly facilitate taxonomic classification. Finally,
iii. combining the information obtained with external data, such as weather variables, allows novel insights into insect population dynamics.

For the first hypothesis, we analyze the performance of the device, in terms of the quality of the captured data. The behavior of insects and their exit patterns are analyzed. Additionally, since future plans include capturing flapping frequency (also called wing beat frequency, WBF) data, we evaluate how often individuals fly or at least flap their wings inside the device. For the second hypothesis, we evaluate the efficiency of the iNaturalist AI system for taxonomic classification. Since only a proof of concept was performed, no simultaneous monitoring with conventional traps was performed for comparison. However, the level of identification of the system is analyzed for each observation. In addition, the community’s rectifications or ratifications and the accuracy of their reviews are also evaluated. Finally, the third approach explores the potential of real-time insect data. However, an in-depth analysis awaits future research.

Overall, we expect to demonstrate the potential of combining field-based generalist non-lethal monitoring devices with an AI-based curated taxonomic image classification platform to allow for automatic and real-time monitoring of insects, applicable also to future larger-scale monitoring programs, especially in cases where lethal traps are not an option.

## Materials and Methods

The FAIR-Device was primarily designed to replace the collection bottle attached to the outlet of a Malaise trap. These generalist interception traps are made of nets of variable thickness, depending on the target species, arranged in an open tent-like form to intercept flying insects such as Diptera and Hymenoptera. Intercepted insects that climb or fly upwards to escape are funneled to the peak of the trap and eventually find their way into a collector jar with a lethal and preservative liquid. To allow non-lethal trapping, the design of the FAIR-Device’s casing allows insects to exit the device. Additionally, a computer surveillance system monitors changes in the field of view of the camera inside the device. When the camera detects the movement of any entering insect above a certain size, a video recording will be generated until the movement stops. The generated video is subsequently saved to the SD card (solid state disk) of the device. The housing design, electronic components, and the method used to count and identify insect species are described in the following sections.

### Trap system

The trapping system tested to bring insects into the FAIR-Device was a Malaise trap developed by the dipterologist M. Bartak (Czech Agricultural University, Prague) [20] with changes made by Bioform (Bioform, Nürnberg, Germany) [21] (**Figure 1**). The trap had dimensions of 270 cm in length, 100 cm in width, and 175 cm in height. The net material consists of UV-resistant 1-mm double-thread fabric.

**Figure 1.**
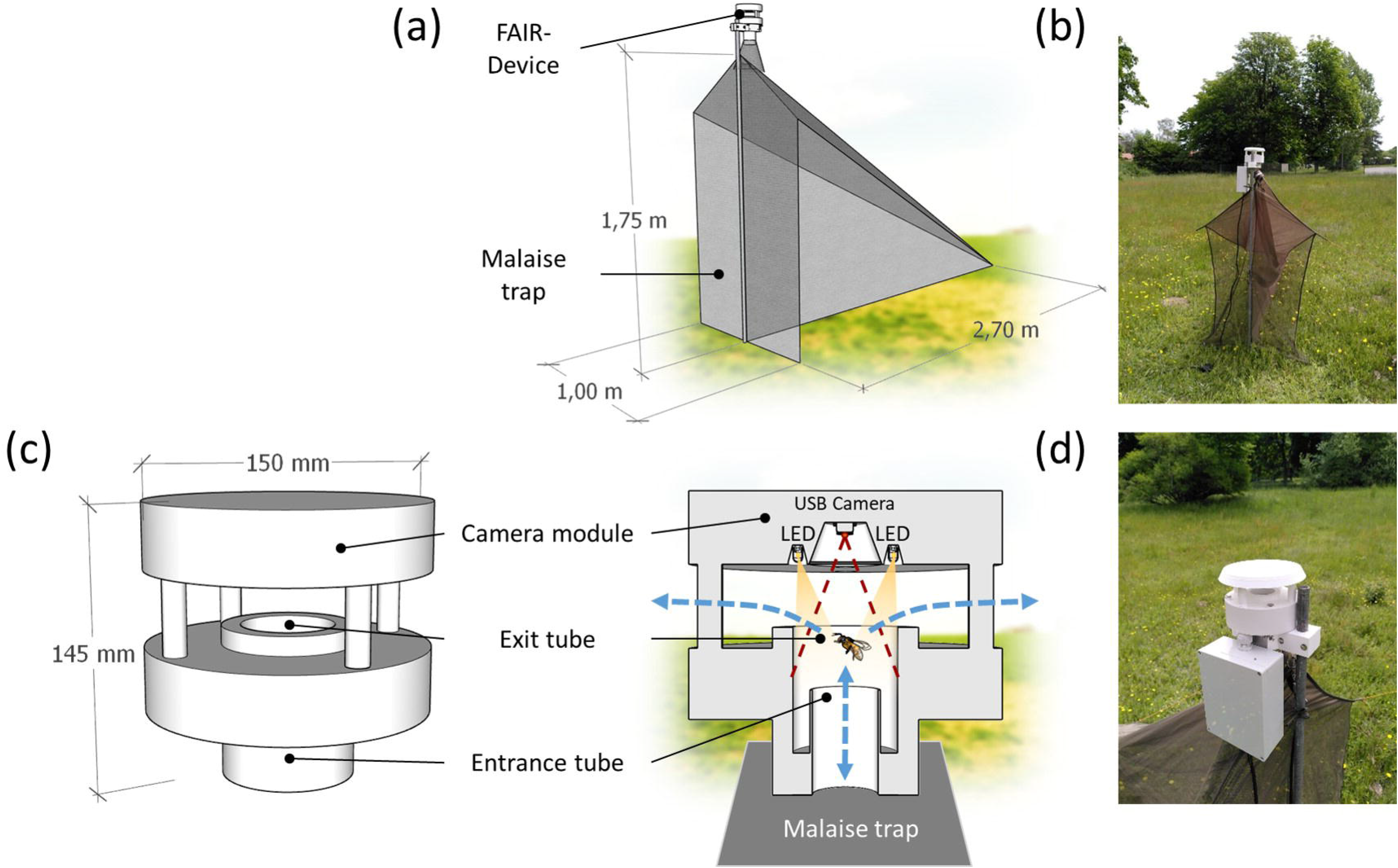
Design of the FAIR-Device. (a) Schematic view of the Malaise trap with FAIR-Device attached on top; (b) FAIR-Device in the field, Summer 2021, Thünen Institute Braunschweig; (c) Close-up of the FAIR-Device with camera module, exit hole, exit, and entrance tubes; (d) Close-up of the device in the field. Images taken by J. Chiavassa.

### Device design and operation principles

Once intercepted by the Malaise trap and funneled upwards, the insects enter the FAIR-Device through a 30 mm wide entrance tube, either by flying or crawling. After reaching the rim of the inlet tube, the insects find themselves in a tube with a larger diameter (50 mm), illuminated from above by four LEDs that provide light to allow image capture and at the same time to attract and induce them to move upwards, towards the device’s exit hole. Once outside, sunlight reduces the attraction effect of the LEDs. The focus of the camera and the LEDs are directed downwards, against the expected direction of insect movement, leaving this configuration with only a few blind spots. The camera lens was calibrated to focus at the exact distance where the rim of the inlet tube is located (74 mm). The movements generated inside the device automatically activate the video recording. A more detailed exploded view of the device will be described in the **Figure 1**.

### Device components and auxiliary systems

The housing components of the device are designed to be fully 3D printed and easy to assemble, which significantly reduces manufacturing costs and allows for dynamic design improvement. The components were 3D-printed with white acrylonitrile styrene acrylate (ASA) filament, a material that is highly resistant to weathering and UV radiation, making it ideal for outdoor applications. The camera module components consisted of a 1.0-megapixel USB camera (ELP720p camera with 45° M7 objective, model USB100W07M-MHV45; Ailipu Technology Co., Ltd., Shenzhen, Guangdong, China) connected to a Single Board Computer (SBC) plus four 5 mm LEDs (334-15/T2C3-2TVC, Everlight Electronics Co., Ltd. NewTaipéi, Taiwan) for lightening the device’s interior. The SBC used was a BeagleBone Black (BeagleBoard.org Foundation, Michigan, USA), a low-cost, SBC with a 1 GHz ARM Cortex-A8 processor, 512MB of DDR3 RAM, and 4GB of eMMC flash storage. It is a community-supported development platform that can run Linux and other operating systems and is commonly used for prototyping and experimentation in fields such as robotics, automation, and the Internet of Things (IoT) [22]. The SBC’s hard drive was a 64 GB SD card (SanDisk Ultra, WesternDigital UK Ltd., Guildford, Surrey, UK).

The support systems of the prototype were the simplest and most reliable possible, as our aim was not a fully autonomous operation of the device in field conditions. The power supply was via connection to the conventional power grid (230 V) with the corresponding conversion to low-voltage direct current (5 V) to power up the SBC. The connection of the SBC to the computer was via a medium-distance WLAN network (EAP110-Outdoor access point, TP-Link Corporation Ltd., Tsim Sha Tsui, Hong Kong).

To obtain real-time weather data, an autonomous weather station was installed at approximately 6 meters distance. The WS501-UMB Smart Weather Sensor (G. Lufft Mess-und Regeltechnik GmbH, Fellbach, Germany) was set to record data on temperature, relative humidity, air pressure, wind direction, wind speed, solar radiation, and precipitation (Lufft rain gauge WTB100) every 15 minutes.

The cost of the device, excluding the Malaise trap, is estimated at approximately 210 Euros.

### Automatic image capture and manual processing

The camera module was operated using a Linux distribution program called MotionEye [23] that was installed on the BeagleBone. The app is written in Python and is a web frontend for the motion daemon, a program mainly developed to manage surveillance cameras. Captured images can be viewed live-streamed and can also be saved as photos or videos (**Figure 2**). It also has widely configurable functionalities for both motion detection to trigger image capture and region of interest masking. MotionEye was configured to record videos instead of images (.mp4 format) since in this way the behavior of the insects inside the device could be observed, as well as knowing with greater precision and less ambiguity the moment of entry and exit of individuals from the device. To minimize the use of the capacity of the SD memory unit, a video resolution of 320×240 pixels was used, considered the lowest possible that still allows obtaining acceptable morphological details of the recorded insects. As it is planned to implement an AI tracking system in the future, it was decided to set the camera to the highest supported recording speed, i.e. 30 fps.

**Figure 2.**
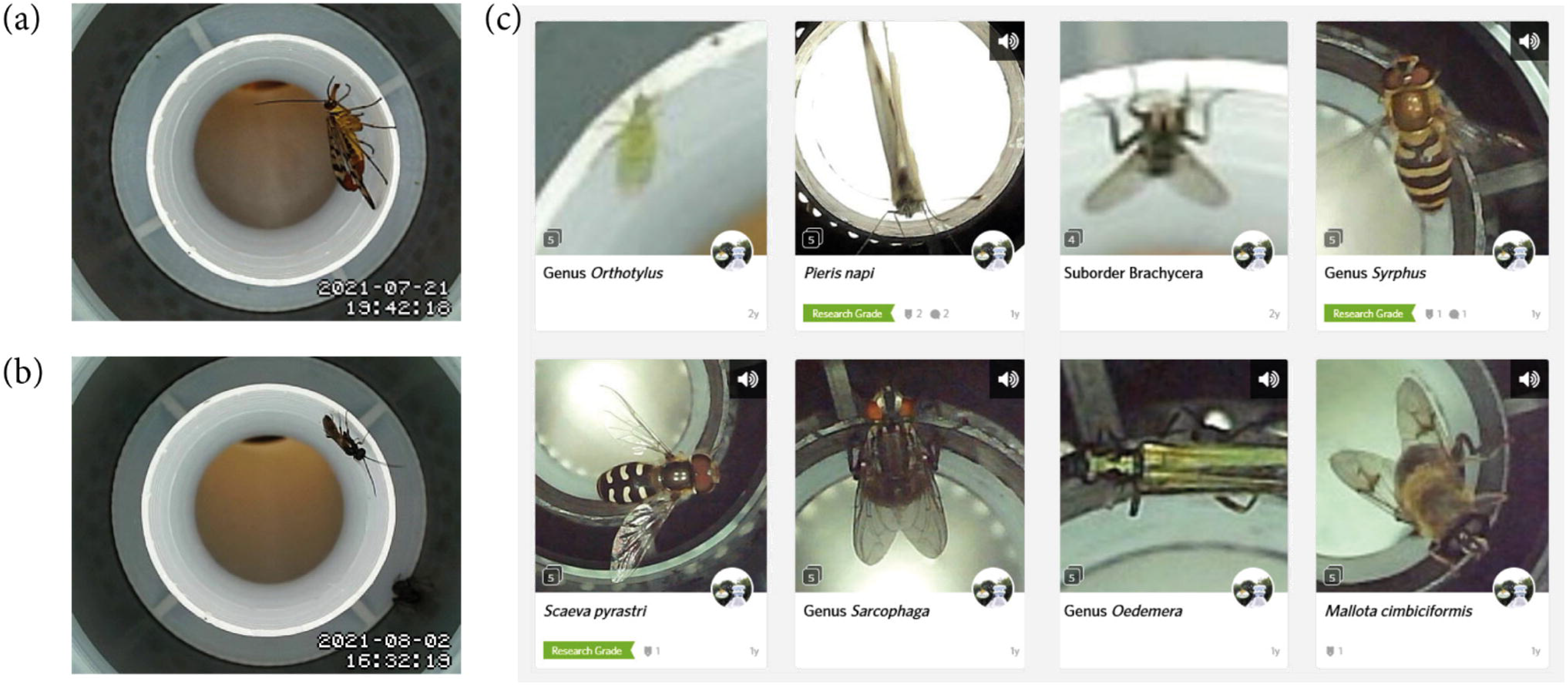
Raw images and image database. (a) A scorpionfly (Panorpa communis L., Mecoptera: Panorpidae) photographed by the camera module on 21 July 2021; (b) an unidentified parasitoid wasp (Hymenoptera: Ichneumonoidea), 2 August 2021; (c) Video frames uploaded to inaturalist.org (screenshot taken 7 July 2023), including suggested taxonomic identifications. Note the “research grade” badges, indicating high certainty in taxonomic identification.

The processing pipeline for the automatic image capture and subsequent database creation for this study is detailed in the flowchart shown in **Figure 3**. The camera’s monitoring system was programmed to detect movement when there were changes in the image of at least 5% and at most 50% of the pixels. This last parameter was set up to avoid, for example, that changes in light (most pixel changes) generate false positives of movement. Likewise, it was established that 120 frames be recorded before and after the movement event (buffer) to determine the exact moment of entry and exit of the individual. If after 20 seconds no new movement was detected, the recording ended, and the video was saved to the SD card with an event start timestamp. Since the test was carried out on the field without constant supervision, we did not evaluate the incidence of false negatives, that is, the entry of an insect into the device without it being detected.

**Figure 3.**
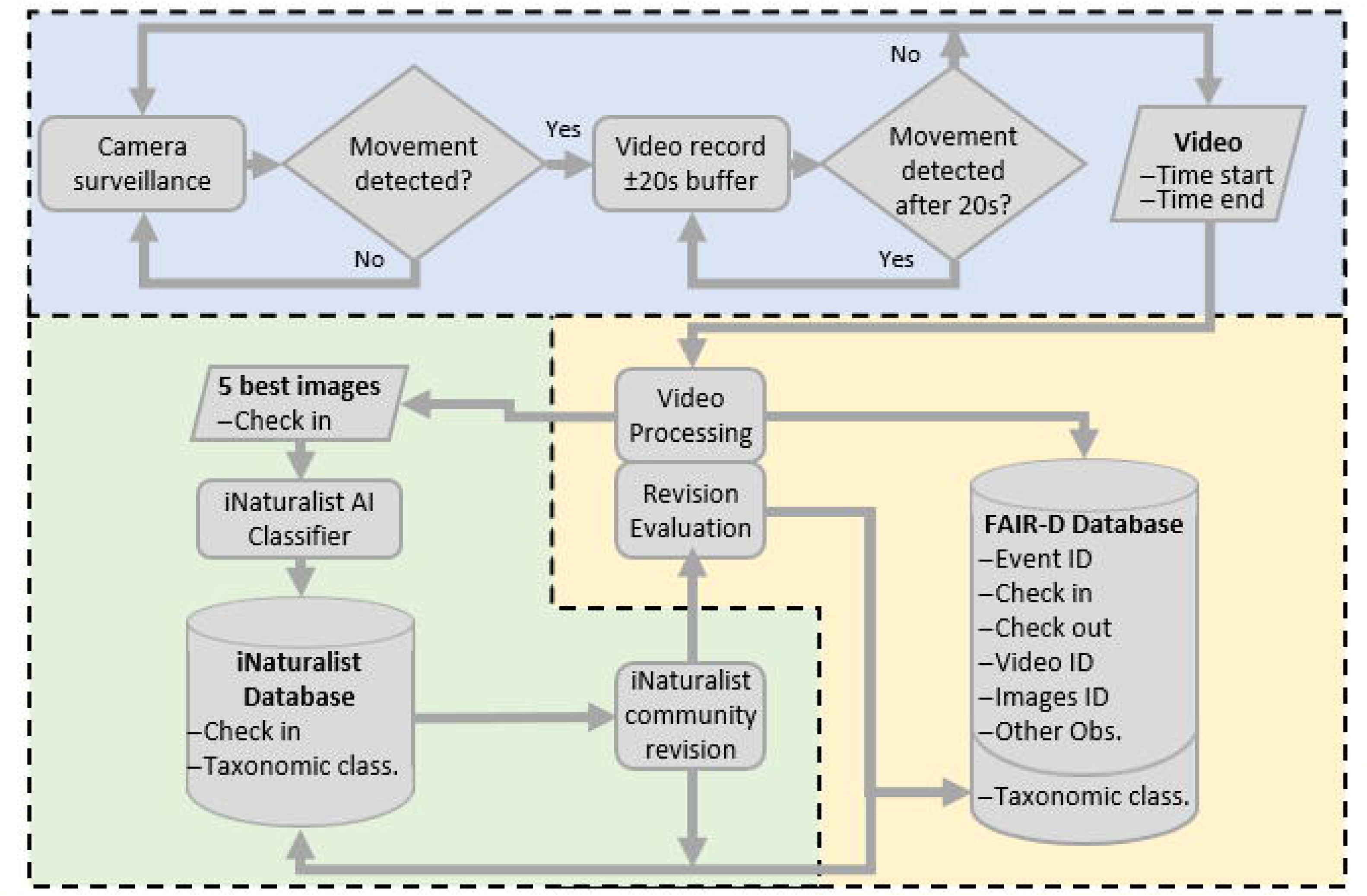
Flowchart of the management of data captured by the device. The blue surface corresponds to the operation of the device in the field, the yellow one to the manual process, and the green one to the process on the iNaturalist platform.

The videos automatically saved by the FAIR-Device were analyzed manually. The first step consisted of eliminating the false positives, i.e., videos where movement was erroneously detected in the absence of an insect. Next, each video containing insects was visually interpreted. Even though two or more individuals may have appeared in the same video, each appearance was treated as a single event. The interpretation process involved annotating relevant information to feed an event database and capturing at least 5 of the best-quality image frames of each individual for subsequent taxonomic classification. Image quality was defined based on images whether the images were (i) not blurred due to the insect’s movement and (ii) overall sharpness. The information of each event registered in the database consisted of the ID of the event, the ID of the video(s) with the same individual, the precise time to the second of its entry (check-in) and exit (check-out), and the ID of the directory to which the extracted images had been saved. Likewise, other observations arising from the video analysis were recorded, such as the insect’s exit direction (again through the entrance or toward the exit) and whether the insect flapped its wings or took flight.

### Taxonomic classification of insects

The taxonomic classification of the insects recorded on video was carried out with the support of the artificial intelligence identification system of the iNaturalist platform. For image data, iNaturalist uses computer vision systems trained on other users’ photos and identifications to provide automated taxonomic ID suggestions [24] that facilitate the identification of the species captured in the observation.

To identify insects using iNaturalist, we uploaded the manually extracted image frames from each individual to the platform. Although automated uploads are not allowed due to a legal requirement on the platform, iNaturalist has developed an application programming interface (API) called VisionAPI that provides direct access to the Computer Vision Model. It allows for automatic image uploads and returns species suggestions based on the iNaturalist computer vision system. However, this feature is not publicly available and only a small number of select individuals/organizations have fee-based access for research or use in other citizen science apps.

Based on the quality of the images and the type of species, the AI system identified individuals at different taxonomic levels displaying one main most probable taxonomic suggestion and eight alternative suggestions. We simulated a completely automatic identification system by taking the main suggestion marked by the AI, even if it did not coincide with our own expert entomological knowledge. If the iNaturalist’s AI indicated that it did not have enough confidence in its suggestions, we classified the observation at the lowest taxonomic level that encompassed all the AÍs taxonomic suggestions. For example, if the system suggested different species belonging to the order Diptera and Hymenoptera but did not provide a main suggestion, we manually assigned the insect to the class Insecta. Observations that could not be determined as an insect were discarded.

Each classification was then made available to the iNaturalist community for ratification, rectification, and/or brought to the most accurate possible taxonomic level. iNaturalist separates observations into two categories: *Casual* and *Research Grade*. In *Research Grade* observations more than 2/3 of the voters agree on the identification of an observation. *Research Grade* observations are considered high-quality and are shared in a scientific database that is free to use for research purposes [25]. As a ground truth test to evaluate the global accuracy of the classifications, we manually reviewed observations subsequently reviewed by the iNaturalist community and accepted or rejected them as appropriate. These data were taken as the definitive classification in our project database.

Identifications were recorded in our database using class, order, suborder, superfamily, family, subfamily, genus, and species. Other intermediate minor ranks such as tribe, subtribe, subgenus, and subspecies were disregarded. While we used the iNaturalist AI system for all classifications, only 39% of them were made public for analysis by the platform community. This was to avoid excessive repetition of detected insect species while ensuring that each species was represented at least once.

### Study site

The study was carried out near the city of Braunschweig, located in the East of the state of Niedersachsen (Lower Saxony), Germany. The device was installed in the test field of the Thünen Institute of Agricultural Technology (52°17’27.6’N 10°26’18.3’E). Local vegetation is a naturalized grassland of 0.4 hectares, surrounded by small broadleaf forests and crop fields.

### Field tests

Field tests were conducted using a single device in July and August 2021 (see **Table S1** in the **Supplementary Materials**). The housing design, camera module, and software operation were tested over a period of 26 days and 300 hours of effective operation. A subperiod of 10 days was selected (between 24^th^ July and 3^rd^ August) over which the device had a continuous operation time. This period is used here to analyze the data in combination with climatic information. During these 10 days, there were N=125 observations with operation hours from 6:00 a.m. to 7 p.m. (CST Time Zone).

### Statistical Analyses

Data were analyzed using R 4.2.3 (2023-03-15 ucrt -- “Shortstop Beagle”) operated via RStudio (2023.06.0 Build 421, © 2009-2023 Posit Software, PBC). Insect counts were analyzed at a range of temporal scales: To analyze daily abundance changes, abundances were summed for each hour (0-24h).

Insect abundances vs. date and stay duration inside the trap were analyzed using a generalized additive model (R package mgcv [26]) with a negative binomial family with either date or log-transformed stay duration as an explanatory variable. Log transformation of stay duration was necessary to account for the highly unequal spread of values in x direction (leverage). The smooth term in the generalized additive model was a penalized thin-plate spline (“ts” in the gam call [27]). The optimal degrees of freedom were estimated using generalized cross validation (option “select=T” in the call to gam).

Responses of insect abundances to temperature and wind speed were analyzed using generalized linear models with negative binomial errors, implemented in the glm.nb function in the MASS library in R [28]. The explanatory variables were check-in date and time, air temperature, and wind speed, all modeled using a B-spline basis of order 4.

Insect order composition over time (dates) was analyzed using multinomial models, implemented in the multinom() function in the nnet package in R (Venables & Ripley 2002). The check-in times of insects in the trap were used; to run the model, these times were transformed to numeric and later back-transformed to dates again. The explanatory variable was the numeric check-in time. A B-spline of order 3 was used to account for non-linearity.

## Results

### Device design performance

Throughout the test period, 942 video files (24.8 GB of data) were recorded, out of which 431 insect individuals were manually registered, with about 16 individuals per day. Despite relatively low resolutions to avoid overloading the SBC, the resulting images were clear enough to distinguish the shapes, textures, and colors of the insects (**Figure 2** (a)). By configuring the camera to record at its maximum possible speed (30 fps), the videos had a sufficient number of frames to produce sharp images in most cases, even when capturing insects in constant motion.

The MotionEye application also met the specifications of properly detecting the entering insects and in most cases fully recording the event (89,7% of the cases). However, some difficulties were encountered with the generation of false positives due to the limitations of the pixel-change-based motion detection system. Despite different settings, about 15% of the total recorded videos were false motion detections. They were usually triggered by sudden changes in light and their occurrence could not be completely prevented. Over the course of the operation period, insect abundance per hour and day fluctuated around 0-5 individuals (2.6 ± 0.2; **Figure 4a**). The penalized thin-plate spline function was of order 3.7 (Chi²=34.9, P<0.005, N=163).

**Figure 4.**
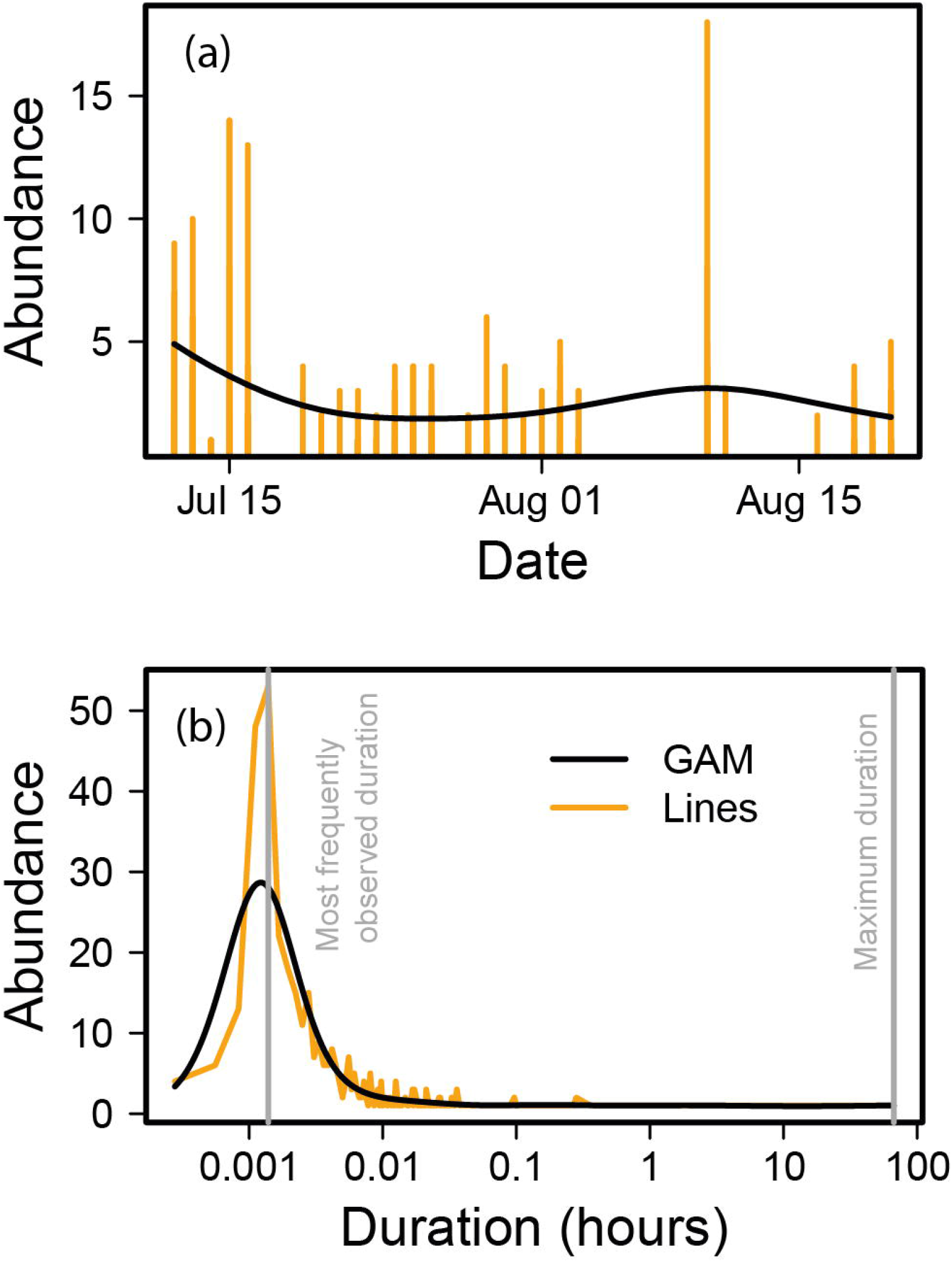
(a) Mean insect abundance per hour per day throughout the operating period of the trap in 2021; (b) average duration of stay of insects inside the traps (in hours). The orange line connects the observations, the smooth black line is a generalized additive model fit. The grey vertical lines indicate the most frequently observed and the maximum duration.

**Figure 5.**
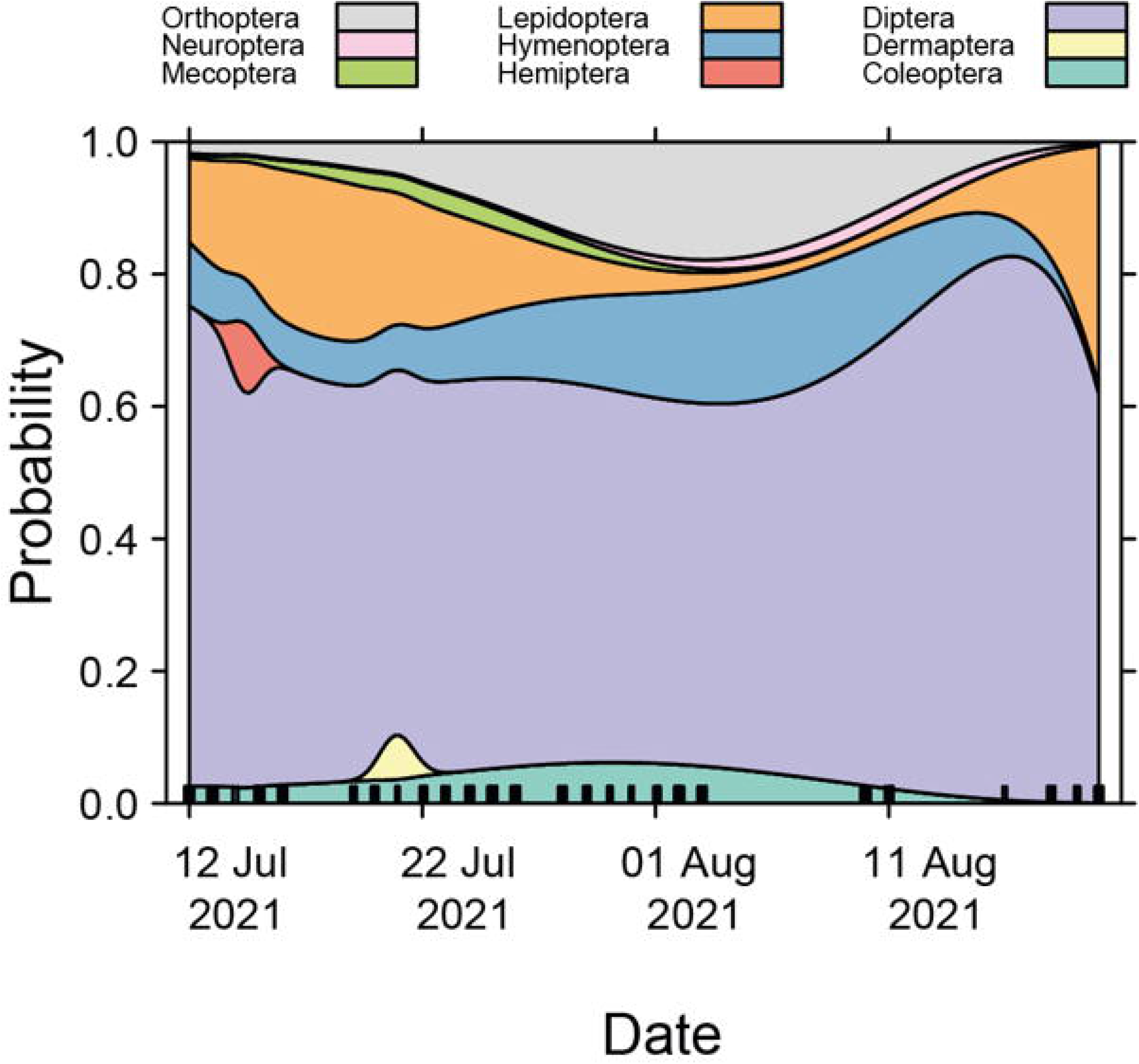
Insect order composition over time. The graph shows predictions from a multinomial model fit using the nnet package in R.

Although the behavior of each insect varied, they all eventually left the device. However, on some occasions, this took several hours, with the longest stay being 66:49 hours (**Figure 4b**) for an ant queen (*Formica sp.*). The most frequently observed stay duration, however, was exactly 5 seconds (**Figure 4b**). This indicates that most of the insects exited the device in a very short time. The corresponding penalized thin plate spline for abundance against stay duration (in hours) had 7.442 d.f. (Chi²=451.8, P<0.001, N=140). When leaving the device, 82,8% of insects exited as intended, while 6,9% returned to the entrance, and 10,3% of their actions were unrecorded. Additionally, 73% of the insects flew or flapped their wings at least once, usually on their way to the exit. The taxonomic order with the greatest tendency to fly was Diptera with 89% of the cases. There were other orders whose individuals did not fly, even though they had wings, as was the case for Orthoptera, Mecoptera, and Neuroptera.

### Insect taxonomy

On the iNaturalist platform, 163 out of the total classifications were made public (**Figure 2c**), which is approximately 39%. The reason for uploading only this small portion of the total observations was to reduce many repeated entries for the same species and thus get a better quality review. The aim was to have at least one published observation for every species. There were two cases in which the AI system could not determine whether the taxon had been an insect or not, proposing also organisms belonging to other kingdoms. These observations were discarded and were not counted as part of the detected individuals. Of the total 163 observations made public in iNaturalist, only 89 (54.6%) were reviewed by the community. The level of review rate by the community can be considered as medium to low. Of the reviewed observations, almost 26% reached the *Research Grade* identification level. The classification made by the iNaturalist AI was in some cases approved by the community (23,9% of the published observations), and in others corrected, in which case it could be classified in a higher taxonomy level (14,1%), at a lower level (8,6%) or corrected to another different taxonomy (8,0%). Regarding the ground truth test for evaluating the global accuracy of the classifications, of 89 observations, 87 were approved by us (97.8%), only one review was rejected (1,1%) and one observation remained undetermined (1,1%).

Most of the insects classified were Diptera (63%), followed by Lepidoptera (13.5%) and Hymenoptera (10%). The B-spline with df=3 for Date in a corresponding multinomial model had Chi²=90.09, df=24, P<0.001. Within the Diptera order, about 60% were fly species that could be directly related to animal production activities. This abundance is likely to be due to the existing pig and milk production facilities near the study site.

The order of Diptera also had the lowest taxonomic resolution obtained by the iNaturalist AI (**Table 1**). While around 99 percent were recognized at the suborder level, only about 54% were categorized at the superfamily level. The Brachycera suborder, which includes flies, was also one of the most difficult to identify at lower taxonomic levels by the iNaturalist AI and community.

**Table 1.** Identification percentage according to the different taxonomic levels and for the different orders within the study period, determined by iNaturalist’s AI identification system and subsequent community review. The higher the level of taxonomic precision, the percentage of identification tends to be lower (transition from green, higher, to red, lower).

| Order | Suborder | Super-family | Family | Subfamily | Genus | Species |
| --- | --- | --- | --- | --- | --- | --- |
| Diptera | 98,87% | 53,58% | 50,19% | 40,38% | 27,55% | 3,40% |
| Hymenoptera | 97,62% | 95,24% | 85,71% | 64,29% | 54,76% | 14,29% |
| Orthoptera | 100,00% | 100,00% | 100,00% | 96,55% | 93,10% | 24,14% |
| Coleoptera | 100,00% | 100,00% | 100,00% | 100,00% | 92,86% | 71,43% |
| Lepidoptera | 87,72% | 84,21% | 54,39% | 36,84% | 35,09% | 10,53% |
| Mecoptera | 100,00% | 100,00% | 100,00% | 100,00% | 100,00% | 0,00% |
| Other | 75,00% | 62,50% | 62,50% | 62,50% | 50,00% | 37,50% |
| Average | 96,90% | 67,38% | 60,24% | 49,29% | 39,29% | 9,76% |

Because the system identified individuals usually at a different taxonomic level, it is not possible to determine exactly how many different species were identified. Without taking into account the non-identifications for the different taxonomic levels, nine orders, fifty families, and 69 different genera were identified. Supplementary Table S2, Table S3, and **Table S4** show all the unique species identified, at the different taxonomic levels and with their respective relative abundance.

### Response to weather variables

Automated monitoring allows responses of insects to be followed as a function of abiotic covariates such as temperature, rainfall, and wind speed. Generalized linear models with negative binomial errors and B-spline basis smooths (df=4) for date and time, wind speed, and temperature had highly significant smooth terms for all three explanatory variables (**Figure 6a**). Insect abundance per 15 minutes (i.e. the finest possible resolution) increased nonlinearly with air temperature and showed a U-shaped response to wind speed (**Figure 6b**). A detailed overview of trap performance, insect count, rainfall events, temperature, and wind speed is shown in **Figure 7**.

**Figure 6.**
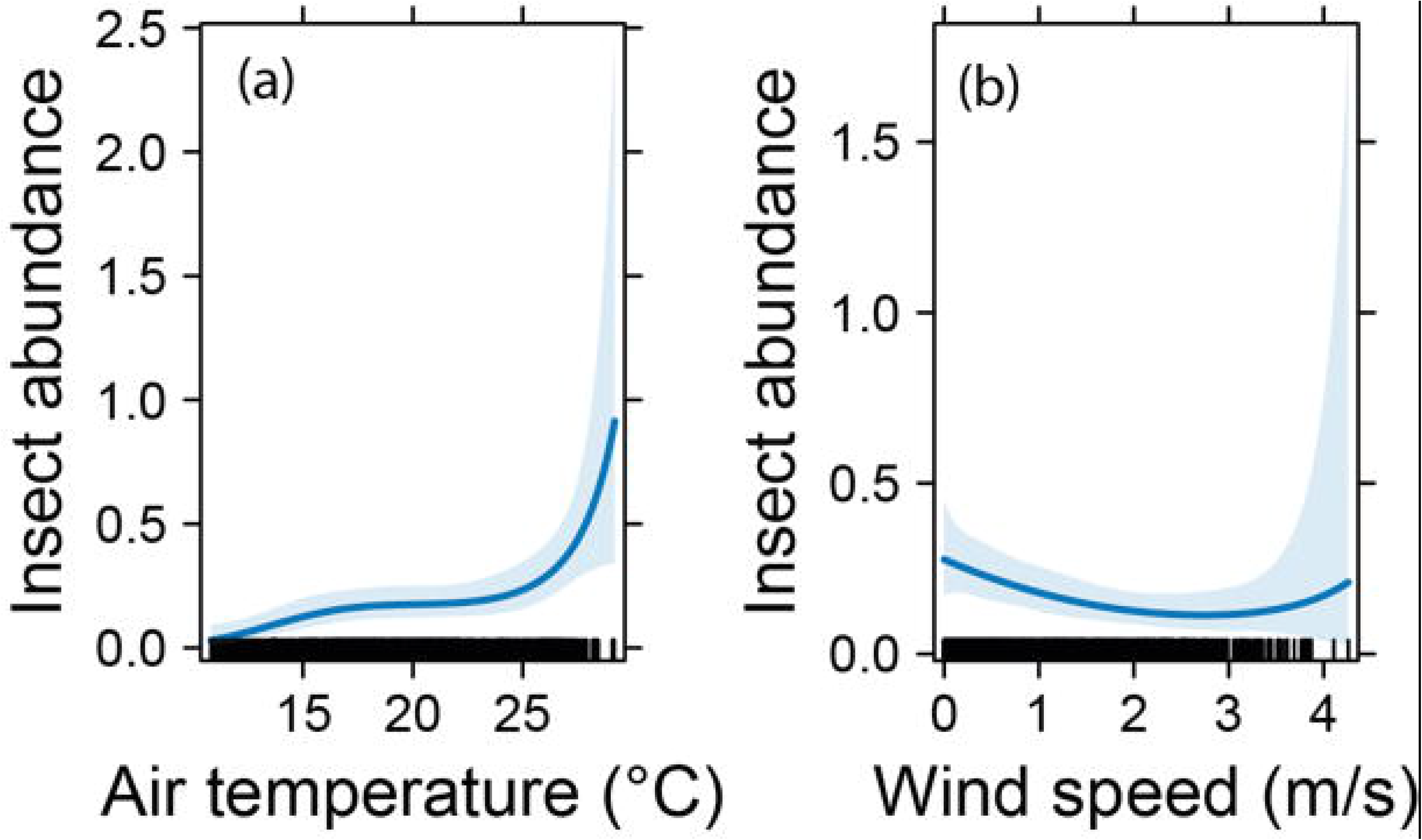
Responses of insect abundance to air temperature (°C) and wind speed (m/s). The lines show back-transformed predictions from generalized linear models, the shaded areas show the 95 percent confidence intervals. Rugs on the x axes show densities of data points (N=3854 time points in total).

**Figure 7.**
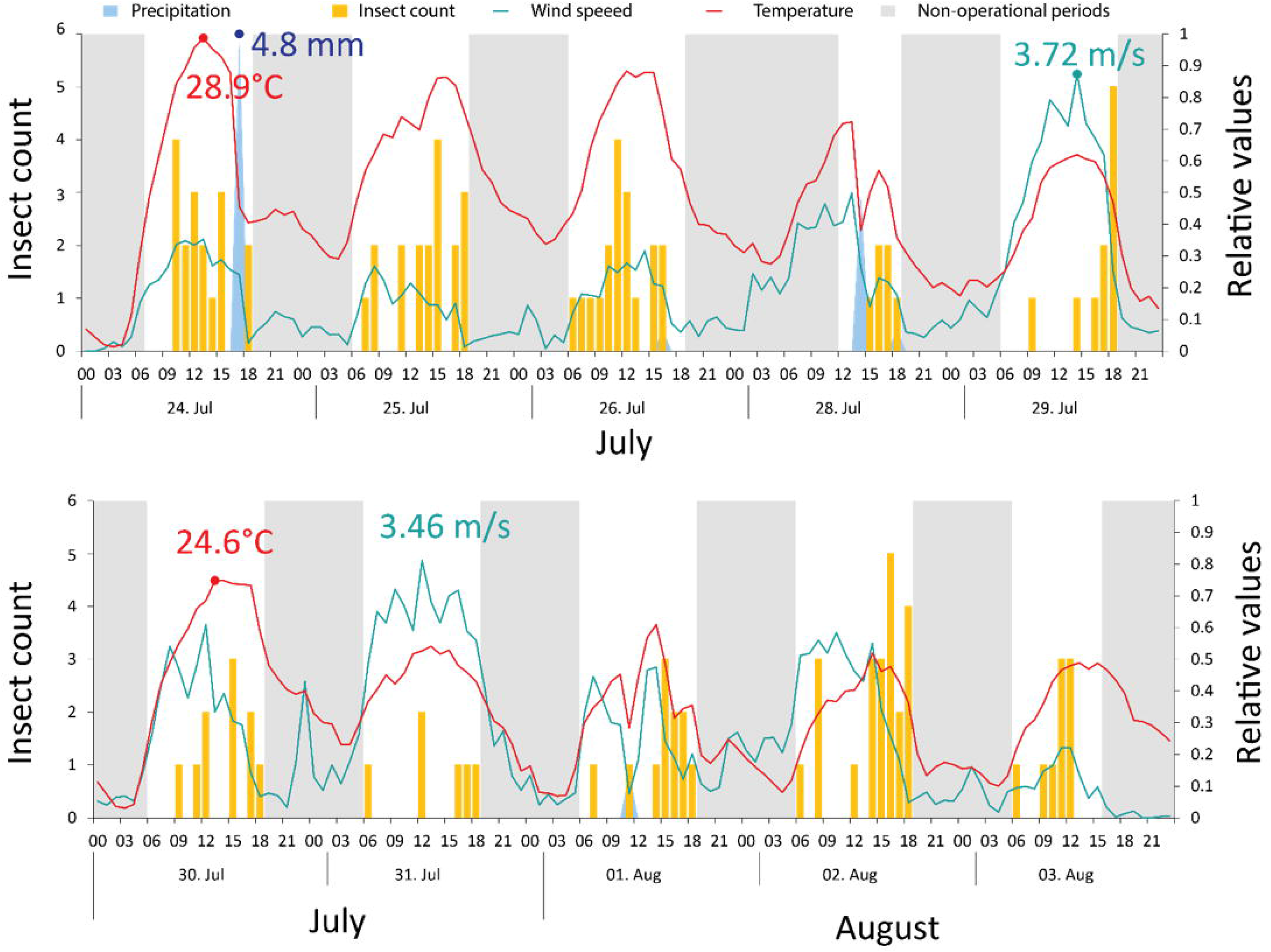
Complex dynamics of insect abundance (yellow), temperature (°C, red), wind speed (m/s, blue-green), and precipitation (mm, blue), including non-operational periods of the trap (grey). Note the relative scale to express all values on a common scale. Relative values are scaled to the maximum of each period, as indicated by the exact values given in the upper and lower panels.

## Discussion

In the present study, we introduce a novel non-lethal Malaise trap system and show a promising avenue to automated insect identification across taxa and orders. Although the system is not yet fully automated, the results obtained are promising, and the taxonomic identification was surprisingly good.

To fully assess taxonomic accuracy, it would have been necessary to catch insects emerging from the devicés exit hole and then let them be identified to species level by taxonomic experts or by DNA metabarcoding approaches for “ground-truthing”. These aspects will be covered in future work.

The FAIR-Device was only operational between July and August, and its operation times did not always cover the full daylight period. Such interruptions occur in any monitoring system and certainly, a more continuous and autonomous operation of the device will be aimed in further developments. Additionally, the device could be supplemented with an intelligent tracking system, where each individual entering could receive a unique ID, avoiding double-counting or problems with the (rarely occurring) lengthy stays inside the trap. As with Malaise traps in general, they measure activity density, and the radius from which insects are attracted is generally unknown. Further work, for example with marking and tracking of individuals, would be needed in order to allow expressing insect observations on a unit area basis (e.g. per square meter).

The availability of qualitative and quantitative data on insects at high temporal resolution opens up fascinating new possibilities. For example, in the case of grassland or cropping systems, responses to management events such as grazing, mowing, or pesticide applications could be studied at unprecedented resolution. This is particularly true for designed experiments, where control and treatment plots could be equipped with FAIR-Devices and simultaneous responses from a multitude of devices could be measured. The resulting data could be used to identify the most effective management strategies for different types of ecosystems, leading to more sustainable and efficient agricultural practices.

A particular strength of equipping a Malaise trap with an automated camera and visual identification system is that the full advantages of Malaise trapping can be used without the need to kill the insects. As such, our approach will be particularly useful for studies in which the conservation of rare species is important and where it would have been difficult to obtain permits for lethal Malaise trapping.

In terms of taxonomic resolution and coverage, while half of the observations remained unrevised in iNaturalist, the taxonomic identification of individuals that were evaluated by the iNaturalist community was excellent. In fact, given the limited resolution of the images, the resulting classifications are outstanding. Thus, platforms such as iNaturalist (or also Observation.org) with their AI algorithms and community-based quality control systems are certainly valuable for the first steps in insect identification from optical monitoring devices. However, the current approach to curating IDs through the platform community may not be scalable in the future. If numerous devices operate simultaneously in the field, sending enormous amounts of data in real-time to be analyzed and reviewed, the percentage of user reviews will likely be significantly lower. On the other hand, these platforms are constantly improving their identification algorithms and computer vision capabilities, which could help mitigate the lack of human supervision. Additionally, to emulate a classification system such as iNaturalist or Observation.org, large image databases for Central European insects taken under field conditions will be needed, combined with efficient classification algorithms (which is outside the scope of the present study).

### Outlook

Many future modifications of the FAIR-Device can be thought of. For example, our imaging head could also be mounted on other types of traps such as vane traps or pitfall traps. Additionally, the device could be equipped with a wing beat frequency module to record the flying events inside the device. This could increase identification precision, especially for those insects with very short stay durations inside the trap. For night-time trapping, the device could also be turned into a light trap by adding blacklight lamps. Overall, the FAIR-Device is certainly only a first step, but we believe it will offer transformative opportunities for future biodiversity monitoring.

### Conclusions

We have presented a device with a design and electronics that allows for obtaining quality time-stamped videos from various species of insects without causing their death. The combination with the iNaturalist platform, thanks to the AI recognition functions and subsequent corrections made by its community, permitted this data to be converted into detailed information about the insect biodiversity of a given site. While there are aspects that still need to be improved or developed, the proposed device has shown promising results for cost-effective, non-lethal, and high temporal and spatial resolution insect biodiversity monitoring using e-traps combined with nature identification platforms.

We believe that automated systems capable of monitoring insects in real-time and 24/7 would open new doors for studying insect populations in ecological research and agricultural production. E-traps arranged as individual nodes to form networks of devices would enable a highly accurate understanding of the spatial and temporal population dynamics of various insect species. Real-time information on pest outbreaks, the effectiveness of control or nature conservation measures, and population data on beneficial or endangered insects could be provided by a network of monitoring devices. Local networks could become nodes of larger networks, eventually reaching global coverage [29]. This would allow both agricultural and conservation research communities to benefit from the same data and indicators, leading to better-informed choices for the long-term economic and environmental viability of both productive and natural systems.

## Author Contributions

Conceptualization, J.A.Ch. and M.K.; methodology, J.A.Ch.; software, J.A.Ch.; validation, J.A.Ch.; statistical analysis, C.S.; data curation, J.A.Ch.; writing—original draft preparation, J.A.Ch.; writing—review and editing, J.A.Ch., M.K., C.S., A.K., P.N., S.W.; visualization, J.A.Ch., C.S.; supervision, J.A.Ch., C.S., A.K., P.N., S.W.; project administration, M.K..; funding acquisition, M.K. All authors have read and agreed to the published version of the manuscript.

## Funding

The presented study was part of the joint project “Monitoring of biodiversity in agricultural landscapes” (MonViA) that has been funded by the German Federal Ministry of Food and Agriculture.

## Acknowledgments

The authors would like to thank Ulf Prüße for the many detailed reviews and advice, Cristina Umstätter for her advice and suggestions, and Alexander Huf for his technical support.

## Conflicts of Interest

The authors declare no conflict of interest.

## Supplementary Materials

**Table S1.** Operating time (CST) of the FAIR-Device in July (top) and August (bottom) 2021.

| Operation | Jul 21 |  |  |  |  |  |  |  |  |  |  |  |  |  |  |  |  | Total |
| --- | --- | --- | --- | --- | --- | --- | --- | --- | --- | --- | --- | --- | --- | --- | --- | --- | --- | --- |
| Days | 12 | 13 | 14 | 15 | 16 | 19 | 20 | 21 | 22 | 23 | 24 | 25 | 26 | 28 | 29 | 30 | 31 | 17 days |
| Start | 13:00 | 7:00 | 7:30 | 18:00 | 0:00 | 20:00 | 0:00 | 19:00 | 0:00 | 0:00 | 7:00 | 6:00 | 6:00 | 12:00 | 6:00 | 6:00 | 6:00 | - |
| Finish | 21:00 | 21:00 | 9:30 | 23:59 | 18:00 | 23:59 | 18:00 | 23:59 | 23:59 | 18:30 | 19:00 | 19:00 | 19:00 | 19:00 | 19:00 | 19:00 | 19:00 | - |
| Effective time | 8:00 | 14:00 | 2:00 | 5:59 | 18:00 | 3:59 | 18:00 | 4:59 | 23:59 | 18:30 | 12:00 | 13:00 | 13:00 | 7:00 | 13:00 | 13:00 | 13:00 | 201:29 |

| Operation | Aug 21 |  |  |  |  |  |  |  |  | Total |
| --- | --- | --- | --- | --- | --- | --- | --- | --- | --- | --- |
| Days | 1 | 2 | 3 | 10 | 11 | 16 | 18 | 19 | 20 | 9 days |
| Start | 6:00 | 6:00 | 6:00 | 15:30 | 0:00 | 12:00 | 18:00 | 15:30 | 0:00 |  |
| Finish | 19:00 | 19:00 | 16:00 | 23:59 | 15:00 | 13:00 | 23:59 | 23:59 | 23:59 |  |
| Effective time | 13:00 | 13:00 | 10:00 | 8:29 | 15:00 | 1:00 | 5:59 | 8:29 | 23:59 | 98:59 |

**Table S2.**
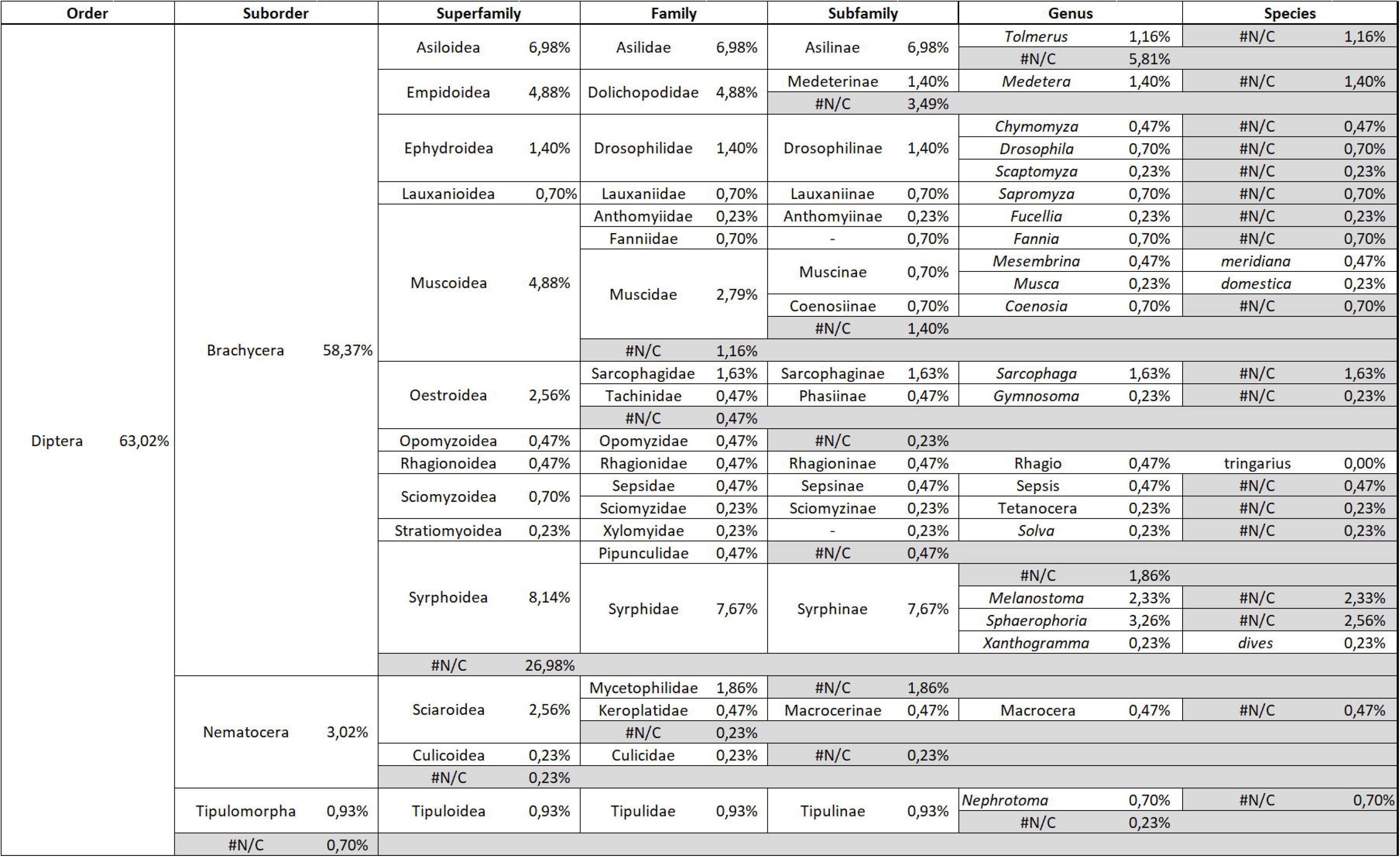
Total unique species identified for Diptera, at the different taxonomic levels and with their respective relative abundance. The abbreviation #N/C stands for “not classified”.

**Table S3.**
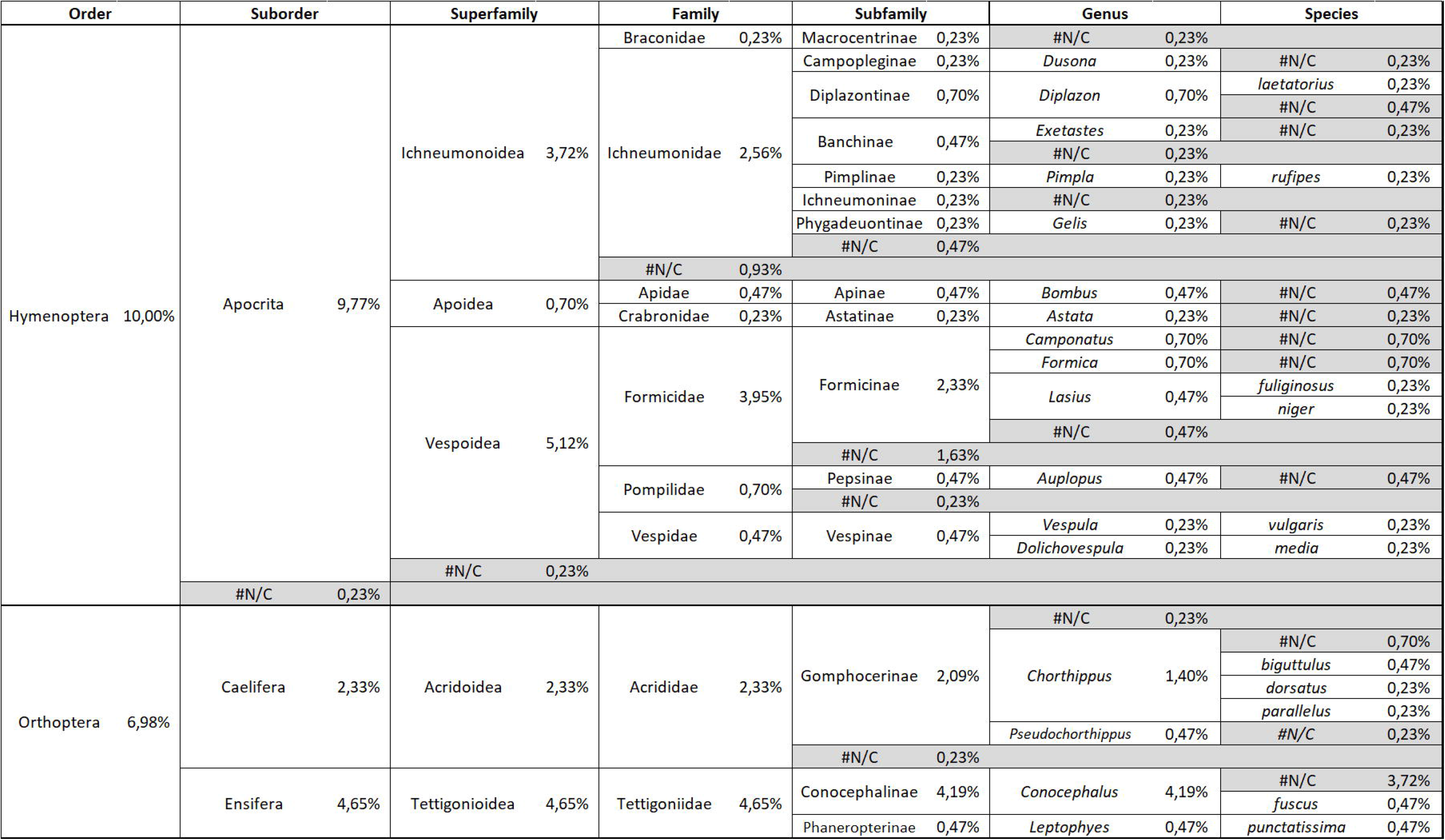
Total unique species identified for orders Hymenoptera and Orthoptera, at the different taxonomic levels and with their respective relative abundance. The abbreviation #N/C stands for “not classified”.

**Table S4.**
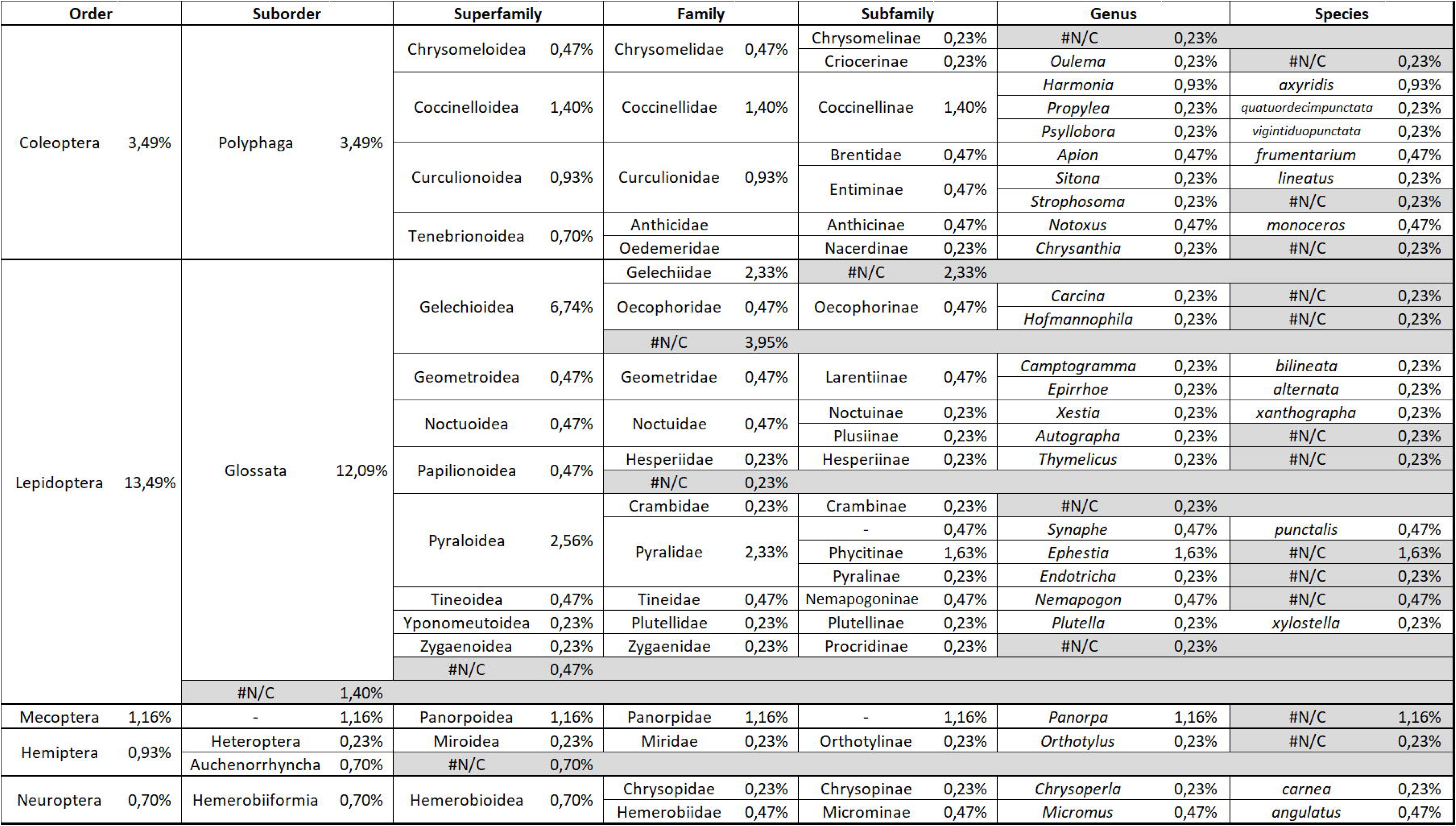
Total unique species identified for orders Coleoptera, Lepidoptera, Mecoptera, Hemiptera, and Neuroptera, at the different taxonomic levels and with their respective relative abundance. The abbreviation #N/C stands for “not classified”.

## Notes

### Competing Interest Statement

The authors have declared no competing interest.

### Summary of Updates

Improved image quality. Added missing authors Orcid IDs.

## References

1. Hallmann CA, Sorg M, Jongejans E, Siepel H, Hofland N, Schwan H, et al. More than 75 percent decline over 27 years in total flying insect biomass in protected areas. PLoS ONE. 2017;12:e0185809. doi:10.1371/journal.pone.0185809.

2. Ceballos G, Ehrlich PR, Dirzo R. Biological annihilation via the ongoing sixth mass extinction signaled by vertebrate population losses and declines. Proc Natl Acad Sci U S A. 2017;114:E6089–E6096. doi:10.1073/pnas.1704949114.

3. Høye TT, Ärje J, Bjerge K, Hansen OLP, Iosifidis A, Leese F, et al. Deep learning and computer vision will transform entomology. Proc Natl Acad Sci U S A 2021. doi:10.1073/pnas.2002545117.

4. Preti M, Verheggen F, Angeli S. Insect pest monitoring with camera-equipped traps: strengths and limitations. J Pest Sci. 2021;94:203–17. doi:10.1007/s10340-020-01309-4.

5. Porter JH, Nagy E, Kratz TK, Hanson P, Collins SL, Arzberger P. New Eyes on the World: Advanced Sensors for Ecology. BioScience. 2009;59:385–97. doi:10.1525/bio.2009.59.5.6.

6. Li X, Zhao N, Jin R, Liu S, Sun X, Wen X, et al. Internet of Things to network smart devices for ecosystem monitoring. Science Bulletin. 2019;64:1234–45. doi:10.1016/j.scib.2019.07.004.

7. Cardim Ferreira Lima M, Damascena de Almeida Leandro ME, Valero C, Pereira Coronel LC, Gonçalves Bazzo CO. Automatic Detection and Monitoring of Insect Pests—A Review. Agriculture. 2020;10:161. doi:10.3390/agriculture10050161.

8. Sciarretta A, Calabrese P. Development of Automated Devices for the Monitoring of Insect Pests. Curr Agri Res Jour. 2019;7:19–25. doi:10.12944/CARJ.7.1.03.

9. Ascolese R, Gargiulo S, Pace R, Nappa P, Griffo R, Nugnes F, Bernardo U. EΛtraps: A valuable monitoring tool to be improved. EPPO Bulletin. 2022;52:175–84. doi:10.1111/epp.12838.

10. Rydhmer K, Bick E, Still L, Strand A, Luciano R, Helmreich S, et al. Automating insect monitoring using unsupervised near-infrared sensors. Sci Rep. 2022;12:2603. doi:10.1038/s41598-022-06439-6.

11. Sittinger M, Uhler J, Pink M, Herz A. Insect Detect: An open-source DIY camera trap for automated insect monitoring. bioRxiv 2023. doi:10.1101/2023.12.05.570242.

12. Sittinger M. Insect Detect Docs. 31.01.2024. https://maxsitt.github.io/insect-detect-docs/. Accessed 6 Feb 2024.

13. Bjerge K, Nielsen JB, Sepstrup MV, Helsing-Nielsen F, Høye TT. An Automated Light Trap to Monitor Moths (Lepidoptera) Using Computer Vision-Based Tracking and Deep Learning. Sensors 2021. doi:10.3390/s21020343.

14. Yao Q, Feng J, Tang J, Xu W, Zhu X, Yang B, et al. Development of an automatic monitoring system for rice light-trap pests based on machine vision. Journal of Integrative Agriculture. 2020;19:2500–13. doi:10.1016/S2095-3119(20)63168-9.

15. van Horn G, Cole E, Beery S, Wilber K, Belongie S, Aodha OM. Benchmarking Representation Learning for Natural World Image Collections; 30.03.2021.

16. iNaturalist - Apps on Google Play. 23.10.2023. https://play.google.com/store/apps/details?id=org.observation.obsidentify&hl=en_US&gl=US. Accessed 23 Oct 2023.

17. ObsIdentify - Apps on Google Play. 23.10.2023. https://play.google.com/store/apps/details?id=org.observation.obsidentify&hl=en_US&gl=US. Accessed 23 Oct 2023.

18. App&nbsp, Store. iNaturalist. 23.10.2023. https://apps.apple.com/de/app/inaturalist/id421397028. Accessed 23 Oct 2023.

19. App&nbsp, Store. ObsIdentify. 23.10.2023. https://apps.apple.com/de/app/obsidentify/id1464543488. Accessed 23 Oct 2023.

20. ResearchGate. Miroslav BARTAK | Professor | Professor | Czech University of Life Sciences Prague, Prague | CULS | Faculty of Agrobiology, Food and Natural Resources | Research profile. 23.10.2023. https://www.researchgate.net/profile/Miroslav-Bartak. Accessed 23 Oct 2023.

21. Bioform. Malaise-Falle bioform nach Bartak. 17.10.2023. https://www.bioform.de/shop.php?action=tree&wg=1&pid=706&treeid=225. Accessed 17 Oct 2023.

22. BeagleBoard.org Foundation. BeagleBoard Black. 05/18/2022 10:56:47. https://beagleboard.org/black. Accessed 18 May 2022.

23. Calin Crisan. motionEye: A web frontend for the motion daemon. 07/15/2022 08:27:09. https://github.com/motioneye-project/motioneye. Accessed 15 Jul 2022.

24. iNaturalist. Help · iNaturalist. 06/16/2022 15:41:59. https://www.inaturalist.org/pages/help#general3. Accessed 16 Jun 2022.

25. Boone ME, Basille M. Using iNaturalist to Contribute Your Nature Observations to Science. EDIS. 2019;2019:5. doi:10.32473/edis-uw458-2019.

26. Wood SN. Generalized additive models: An introduction with R. Boca Raton, FL: CRC Press, Taylor & Francis Group; 2017.

27. Wood SN. Thin Plate Regression Splines. Journal of the Royal Statistical Society Series B: Statistical Methodology. 2003;65:95–114. doi:10.1111/1467-9868.00374.

28. Chambers J, Eddy W, Härdle W, Sheather S, Tierney L, Venables WN, Ripley BD. Modern Applied Statistics with S. New York, NY: Springer New York; 2002.

29. Potamitis I, Eliopoulos P, Rigakis I. Automated Remote Insect Surveillance at a Global Scale and the Internet of Things. Robotics. 2017;6:19. doi:10.3390/robotics6030019.

